# DOCKGROUND Membrane Protein-Protein Set

**DOI:** 10.1101/2021.11.04.467360

**Authors:** Ian Kotthoff, Petras J. Kundrotas, Ilya A. Vakser

**Author notes:** <u>Corresponding authors:</u> Ilya A. Vakser and Petras J. Kundrotas, Computational Biology Program, The University of Kansas, 2030 Becker Drive, Lawrence, Kansas 66045, and.

## Abstract

Membrane proteins play essential role in cellular mechanisms. Despite that and the major progress in experimental structure determination, they are still significantly underrepresented in Protein Data Bank. Thus, computational approaches to protein structure determination, which are important in general, are especially valuable in the case of membrane proteins and protein-protein assemblies. Due to a number of reasons, not the least of which is much greater availability of structural data, the main focus of structure prediction techniques has been on soluble proteins. Structure prediction of protein-protein complexes is a well-developed field of study. However, because of the differences in physicochemical environment in the membranes and the spatial constraints of the membranes, the generic protein-protein docking approaches are not optimal for the membrane proteins. Thus, specialized computational methods for docking of the membrane proteins must be developed. Development and benchmarking of such methods requires high-quality datasets of membrane protein-protein complexes. In this study we present a new dataset of 456 non-redundant alpha helical binary complexes. The set is significantly larger and more representative than previously developed ones. In the future, this set will become the basis for the development of docking and scoring benchmarks, similar to the ones developed for soluble proteins in the DOCKGROUND resource http://dockground.compbio.ku.edu.

## Introduction

Membrane proteins account for a large part (up to 40%) of the human proteome. These proteins individually and in association with other membrane proteins, perform a wide range of functions, such as transporting nutrients, maintaining electrochemical gradients, cell-cell signaling, and structural support.^1^ Recent advances in cryogenic electron microscopy have made it possible to determine the structure of increasingly large number of membrane proteins.^2^ However, they are still significantly underrepresented among structures in the Protein Data Bank.^2,3^ Experimental determination of the 3D structures of protein-protein complexes, in general, is more difficult than that of the individual proteins, compounding the difficulty of determining protein structures in the membrane. Thus, computational methods for prediction of protein-protein complexes (protein docking) are essential for structural characterization of protein-protein interactions in the membranes. The membrane environment constrains the structure of protein-protein interactions by limiting protein insertion angles and depths.^4,5^ Thus, the dimensionality of the docking space in membranes is less than that for the soluble protein-protein complexes. However, the recognition factors in membrane proteins are smaller in scale than in the soluble protein-protein complexes (where coarse-grained representation determined by the global fold often suffices for a meaningful prediction) and thus require atomic-level prediction accuracy.^6,7^

Because of the combination of structural and physicochemical characteristics of the membrane proteins that distinguish them from the soluble ones and the specifics of the membrane environment, docking methodologies developed for the soluble proteins are not optimal for the membrane proteins.^8^ Thus, specialized computational methods for docking of the membrane proteins have to be developed. In order to accomplish that, one needs high-quality datasets of membrane protein-protein complexes, necessary for the development and benchmarking of such methods. The sets have to be large enough to ensure statistical reliability of the results. Existing sets of membrane protein-protein complexes, contain a relatively small numbers of entries.^4^ Koukos et al. describe a complex set of 37 transmembrane targets.^9^ The Memdock benchmark consists of 65 target alpha helical transmembrane complexes.^8^ In this paper we present a new dataset of 456 non-redundant alpha helical binary complexes, as the foundation for the future development of the comprehensive resource for structural studies of membrane protein-protein complexes.

## Results and Discussion

### Generation of Dataset

Initial PDB biounit structures used in this study were downloaded from the Orientation of Proteins in Membrane’s alpha helical transmembrane database.^10^ This database contains both the structure of the protein and the computationally determined membrane. At the time of retrieval (October 2019), the dataset contained 4,359 alpha helical and 530 beta-barrel protein structures. Beta-barrel membrane proteins are found almost exclusively in Gram-negative bacteria, mitochondria and chloroplasts.^11^ Because of that, the number of such structures is relatively small. Thus, we restricted our set to alpha-helical proteins only. After all monomeric proteins were filtered out, 3,359 entries remained. They were further split by the protein chain. To keep only the transmembrane part of the protein, the extramembrane parts of the structures were deleted. All binary complexes formed between any two chains were considered for a potential complex. Thus, one PDB structure could yield several binary complexes. To characterize the interface size, we used FreeSASA^12^ to calculate solvent accessible solvent area (SASA) buried upon protein binding (for that purpose, treating the membrane proteins like the soluble ones). Two chains were considered interacting if their buried SASA was > 250 Å^2^ per chain. This resulted in 7,964 pairwise combinations of the transmembrane segments.

The redundancy was removed at the level of combined tertiary and quaternary structures. For that, we determined all-against-all TM-scores produced by MM-align^13^ (hereafter referred to as MM-score), an offshoot of the structural alignment program align, specifically designed for aligning multi-chain structures. The dataset was clustered by Highly Connected Subgraphs method^14^ with various MM-score cutoffs ranging from 0.4 to 1.0, to find the optimal threshold value maximizing the number of highly connected clusters (where all members of the clusters have MM-score above the threshold). The clustering thresholds were analyzed by the number of resulting clusters and the fraction of singletons (clusters with one element). Figure 1A shows the number of clusters produced at each level. While lower MM-score cutoffs have the benefit of a larger number of complexes in the set, their drawback is allowing greater structural redundancy. To minimize the drawbacks and maximize the benefits, one can assign the optimal MM-score cutoff at the point at which the number of clusters is significantly less than the next highest score (Figure 1A). Another consideration for choosing the clustering threshold is the minimal number of singleton clusters. Figure 1B shows the fraction of such clusters produced with MM-score cutoffs 0.4 - 0.9. Based on these considerations, comparing the distributions in Figures 1A and 1B, we chose the lowest fraction of the singleton clusters as the clustering threshold (MM-score 0.6). Application of this threshold yielded 456 clusters. The largest cluster contained 851 complexes, 153 clusters were singletons and 48 clusters contained two complexes. Representative structures from the clusters for inclusion into final dataset were those with the best structure resolution. If two or more representative structures had the same resolution, the one with the least missing residues was selected. An exemple of a cluster and its representative is shown in Figure 2.

**Figure 1.**
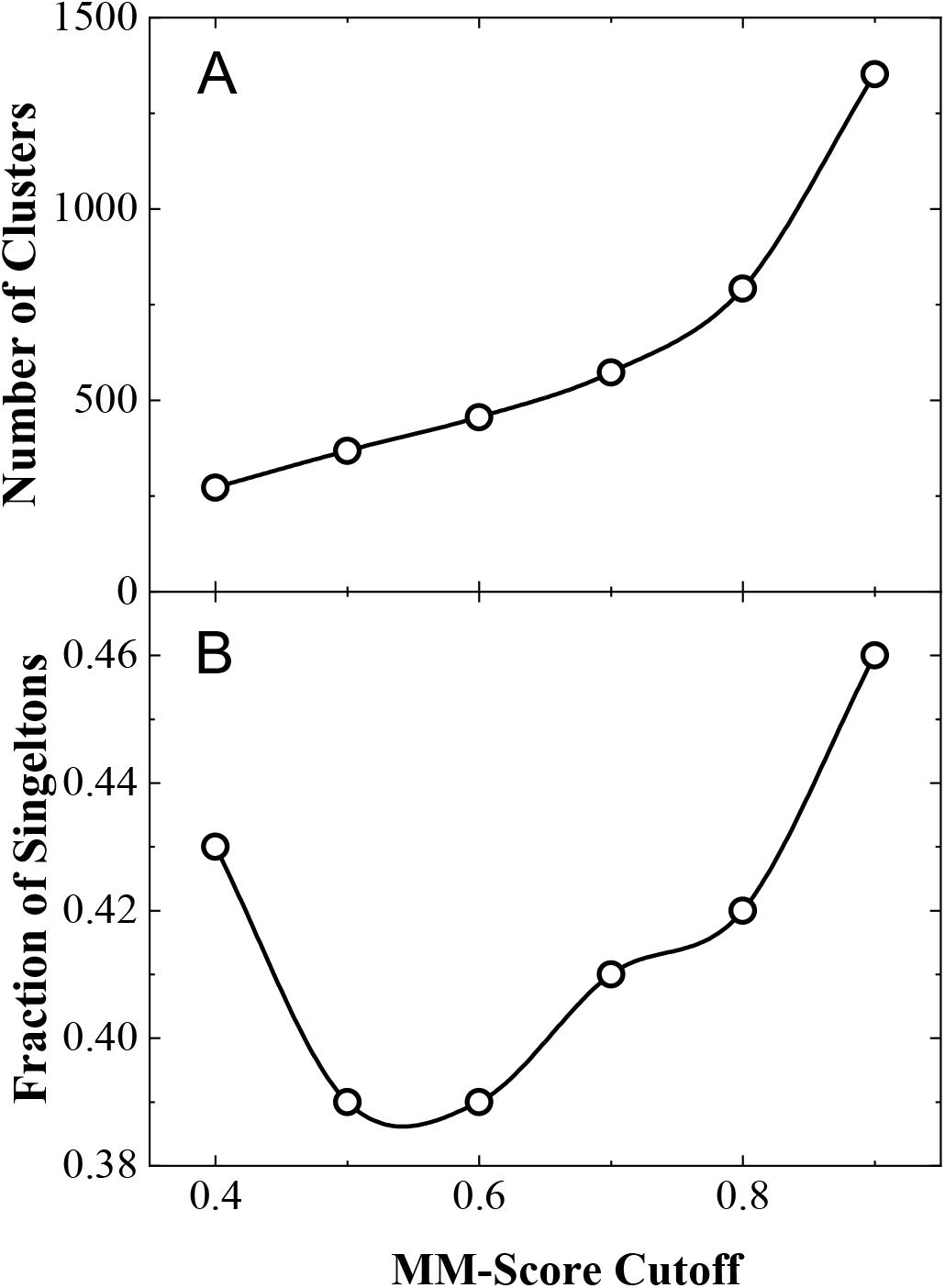
Analysis of structure clusters at different clustering cutoffs. The number of clusters produced at each clustering cutoff (A) grows monotonously, providing no clear indication of the optimal clustering cutoff. The frequency of single complex clusters has a distinct minimum, suggesting MM-score 0.6 as the optimal cutoff.

**Figure 2.**
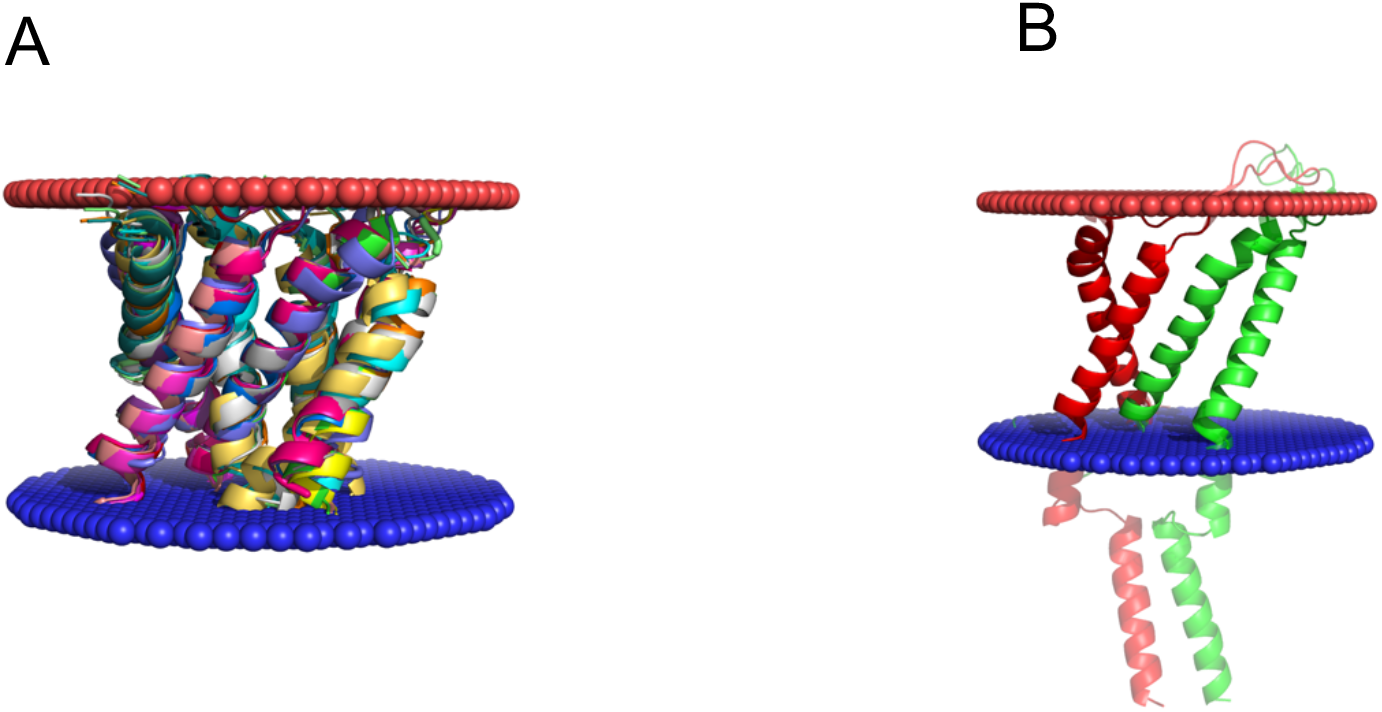
Example of the structure cluster. (A) The cluster consists of 15 binary interactions between transmembrane portions of the protein structures, each having two anti-parallel helices. (B) The cluster’s representative structure contains transmembrane segments of chains A (red) and B (green) from 2wcd. Each of the 15 binary complexes are GO annotated as part of an ion channel. The extra- and intra-cellular sides of the membrane are in red and blue, respectively. The extramembrane parts of the representative complex (not included in the dataset) are blurred for clarity.

### Analysis of Dataset

Membrane environment imposes restrictions on helix insertion angles and depth. We analyzed differences/similarities in the arrangements of helices belonging to the same or different chains. We calculated the angles between all pairs of interacting helices in the final dataset. The distribution of the angles was analyzed separately for the pairs of intra- and inter-chain helices. Angles were calculated between vectors connecting N- and C-termini of a transmembrane helix. The vectors were drawn by performing a linear regression through all C^α^ atoms of the helix. To assign the vector unambiguously, we excluded short helices (those consisting of less than two turns or eight residues). The length of a helix was defined by a continuous stretch of alpha-helical residues as determined by DSSP.^15,16^ With such definition, angles < 90° indicated parallel helices, and those > 90° - the antiparallel ones. We used two alternative distance cutoffs to determine whether a pair of helices is interacting: any C^α^ atom of one helix to any C^α^ atom of the other helix (i) < 6 Å and (ii) < 12 Å (an empirical value based on maximizing docking success rates for soluble proteins^17^). Distributions of the helix-helix angles for both cutoffs (Figure 3) are practically indistinguishable. Thus, here we discuss the results obtained with the 6 Å cutoff only.

**Figure 3.**
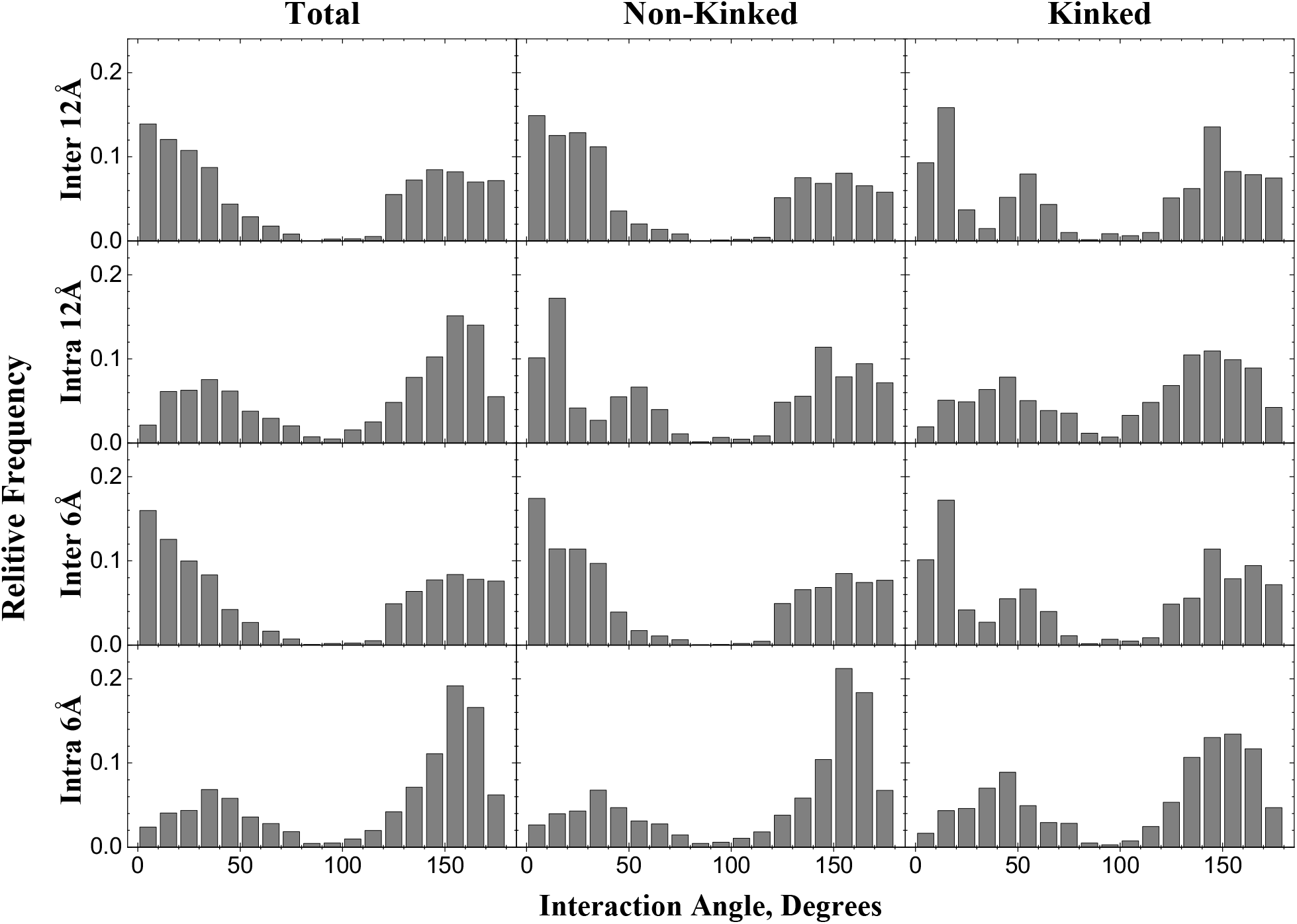
Distribution of angles between interacting intra- and inter-chain helices.

Significant part of the dataset (96 non-redundant entries) contains kinked helices where one or two non-helical residues were present between longer stretches of the alpha-helical residues (≥ 8 residues). For such cases, angles were considered separately between vectors drawn through each part of the kinked helix (Figure 4). This resulted in 1,270 pairs of interacting kinked and 5,725 pairs of non-kinked helices. Distribution of interacting angles for such pairs are shown in Figure 3, along with the distributions of interaction angles between 5,783 of kinked and 16,009 pairs of non-kinked interacting helices, belonging to the same polypeptide chain.

**Figure 4.**
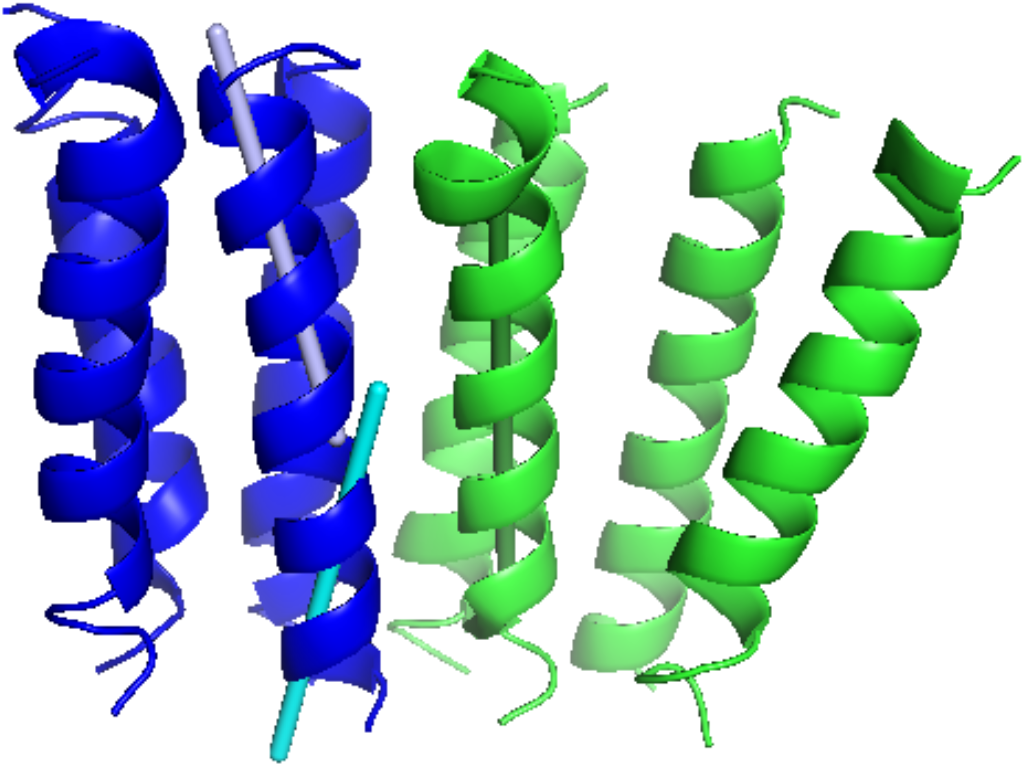
Example of a kinked helix. The transmembrane segments of chain A are in blue, and of chain B are in green (PDB structure 2xq4). Chain A contains a kinked helix with two direction vectors (gray and cyan) used separately in calculation of the interaction angles for that helix.

The inter-chain helix-helix interactions occurred more frequently between parallel helices, whereas the intra-chain interactions were more commonly formed by the anti-parallel helices (sequentially adjacent helices are more likely to interact). Distributions for the kinked helices showed preference for 20-25° kink angles (in ∼50° and ∼140° peaks of the interacting kinked helices distributions). The kinked helices accounted for most interactions angles close to 90° indicating the membrane environment pressure for parallel arrangement of long helices.

The membrane protein-protein set is incorporated in the DOCKGROUND resource for protein recognition studies http://dockground.compbio.ku.edu, in its Bound protein-protein part. The membrane set page (Figure 5) allows download of the entire set, or the individual complexes, along with their visual analysis.

**Figure 5.**
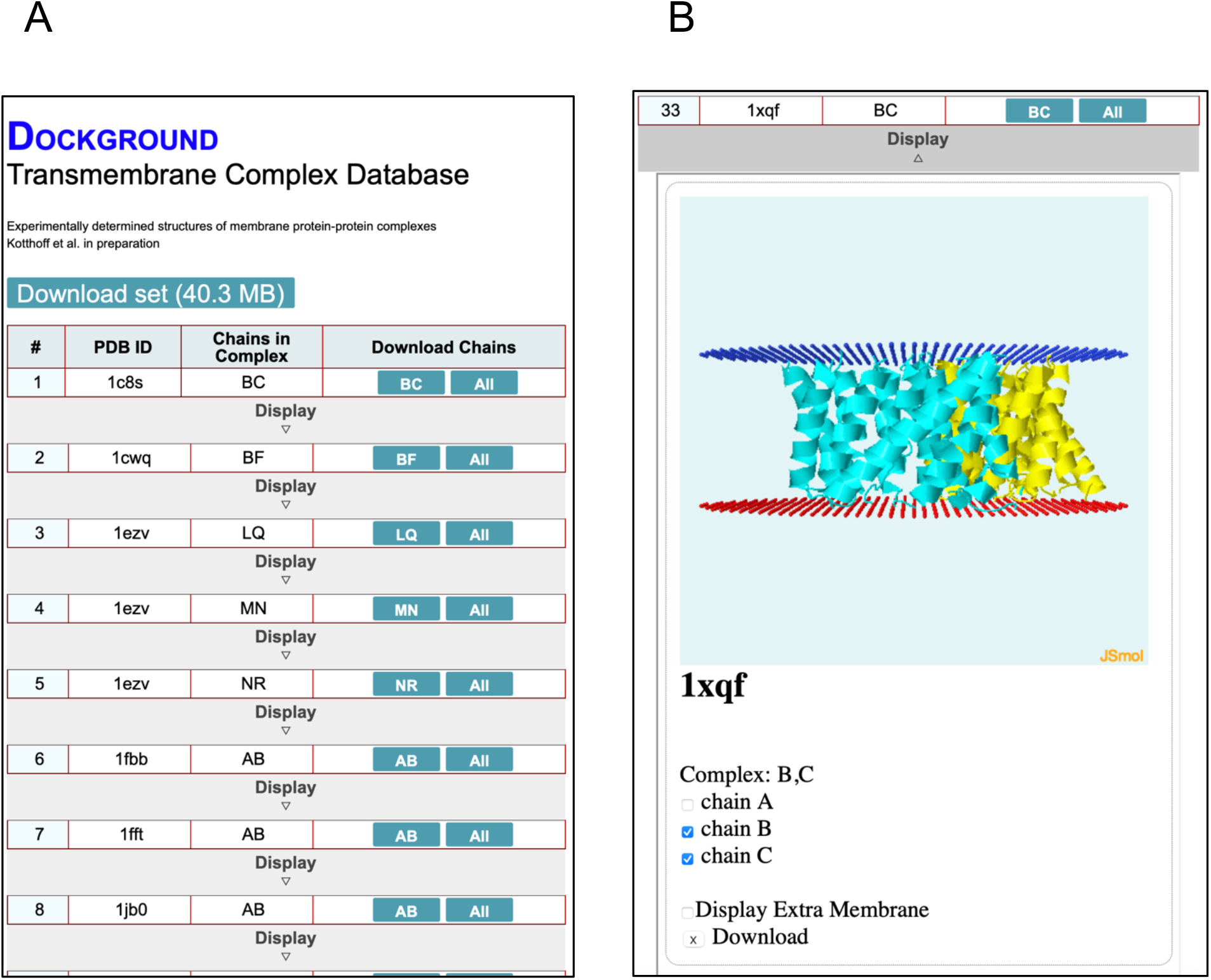
DOCKGROUND webpage for the membrane protein-protein set. (A) The list of complexes for download as a whole or as individual complexes. (B) Visualization of a particular complex.

## Conclusion and Future Work

Membrane proteins play an essential role in cellular mechanisms. Despite that and the major progress in experimental structure determination, they are still significantly underrepresented in Protein Data Bank. While computational approaches to protein structure determination are important in general, they are especially valuable in the case of membrane proteins and protein-protein assemblies. Due to a number of reasons, not the least of which is much greater availability of structural data, the main focus of structure prediction techniques has been on soluble proteins. Structure prediction of protein-protein complexes is a well-developed field of study. However, because of the differences in physicochemical environment in the membranes and the spatial constraints of the membranes, the generic protein-protein docking approaches are not optimal for the membrane proteins. Thus, specialized computational methods for docking of the membrane proteins must be developed. Development and benchmarking of such methods requires high-quality datasets of membrane protein-protein complexes. In this study we present a new dataset of 456 non-redundant alpha helical binary complexes, which is significantly larger and more representative than previously developed sets.

In the future, this set will become the basis for the development of docking and scoring benchmarks, similar to the ones developed for soluble proteins in the DOCKGROUND resource. The sets will contain unbound/modeled structures of the monomers (docking benchmark sets) and docking decoys (scoring benchmark sets) containing correct (near native) and incorrect predictions (decoys) for the development of scoring procedures and training of machine learning approaches.

## Availability

The dataset is available online on the DOCKGROUND resource webpage: http://dockground.compbio.ku.edu.

## Acknowledgments

This study was supported by NIH grant R01GM074255 and NSF grant DBI1917263.

## Notes

### Competing Interest Statement

The authors have declared no competing interest.

